# Trehalose-mediated restoration of Mitf corrects autophagy defects and ameliorates neurodegeneration in a *Drosophila* model of Mucopolysaccharidosis VII

**DOI:** 10.1101/2025.11.10.687576

**Authors:** Apurba Das, Rupak Datta

## Abstract

Mucopolysaccharidosis type VII (MPS VII) is a lysosomal storage disease caused by β-glucuronidase (β-GUS) deficiency. A *Drosophila* model of MPS VII, generated by deleting the fly β-GUS ortholog CG2135, exhibited extensive neurodegeneration and impaired brain autophagy. Here, we investigated the molecular basis of this defect and found that β-GUS deficiency caused marked downregulation of Mitf, the master regulator of autophagy-lysosomal biogenesis, accompanied by reduced expression of its target genes and diminished mTOR activity. Trehalose treatment reactivated Mitf and its downstream autophagy gene network, restored mTOR activity, mitigated neurodegeneration, and significantly improved locomotion in flies. Collectively, our findings identify Mitf dysregulation as a key driver of autophagy impairment in the MPS VII brain and highlight Mitf as a promising therapeutic target.

## Introduction

MPS VII is an autosomal recessive lysosomal storage disease caused by a deficiency of β-glucuronidase (β-GUS), an enzyme essential for the degradation of glycosaminoglycans (GAGs). Neurodegeneration is a prominent clinical manifestation, and patients commonly exhibit intellectual disability, delayed speech, hearing impairment, and restricted movement, affecting nearly 80% of diagnosed individuals. (1,2). Although enzyme replacement therapy (ERT) alleviates systemic symptoms, it fails to correct neurological manifestations due to poor penetration across the blood-brain barrier (3). Gene delivery of β-GUS demonstrated partial improvement in mouse models, however, concerns regarding genomic integration and hepatocarcinoma have limited their translational potential (4–7). This emphasizes the need for a deeper understanding and the development of alternative therapeutic strategies for MPS VII.

To dissect the molecular basis of neuropathology, we utilised a *Drosophila* model of MPS VII (8). CG2135^−/−^ flies exhibited key hallmark neuropathological features including dopaminergic neuronal loss, vacuole formation, retinal degeneration, and progressive decline in climbing ability. Recently, we demonstrated that neurodegeneration in CG2135^−/−^ fly is a consequence of defective autophagy induction and its turnover. The decrease in autophagy related genes Atg1, Atg8a and p62 homolog Ref(2)P led to reduced clearance of damaged mitochondria, and lipofuscins. We showed that inability of non-functional mitochondria to produce sufficient ATP is the underlying reason for neuronal death (9). These neurodegenerative signs may be due to altered lysosomal integrity and impaired transcriptional activation of lysosomal and autophagy-related genes.

Autophagy and lysosomal biogenesis are transcriptionally regulated by the basic helix-loop-helix leucine zipper (bHLH-Zip) transcription factor TFEB (10). In *Drosophila*, its homolog Mitf controls the expression of genes within the CLEAR (**C**oordinated **L**ysosomal **E**xpression and **R**egulation) network, including those required for autophagosome formation and lysosomal activity (11,12). This function is conserved in *Drosophila* (13,14), where Mitf mediated regulation of autophagy is essential for neuronal homeostasis (15,16). Consistently, Mitf knockdown in *Drosophila* causes accumulation of autophagic substrates, while reduced Mitf activity induces neurodegeneration in amyotrophic lateral sclerosis (ALS) fly model (11,17). Given their reliance on constitutive autophagy for getting rid of unwanted proteins and damaged organelles, sustained Mitf suppression is expected to critically impair neuronal clearance capacity, leading to neurodegeneration (18,19). Conversely, modulating Mitf-mediated autophagy promotes lysosome-autophagy biogenesis and mitigates neurodegeneration (20–23). Despite these studies, a key unresolved question is whether altered TFEB/Mitf expression and activity contribute to impaired autophagy, lysosomal dysfunction, and the progression of neurodegeneration in MPS VII.

In this study, we sought to determine whether dysregulation of the TFEB/Mitf pathway underlies the autophagy defects and neurodegeneration associated with MPS VII and whether pharmacological activation of Mitf could restore neuronal homeostasis. Using the Drosophila MPS VII model, we identify Mitf as a previously unrecognized molecular target of β-GUS deficiency and demonstrate that loss of Mitf signaling impairs lysosomal autophagy, thereby driving progressive neurodegeneration. Furthermore, given the ability of trehalose to activate TFEB/Mitf-dependent autophagy in other neurodegenerative models, we evaluated its therapeutic efficacy in MPS VII (24–27). Trehalose restored Mitf activity and markedly ameliorated neuropathological and behavioral defects, including impaired locomotion, lipofuscin accumulation, and neuronal apoptosis. To our knowledge, this is the first study to establish Mitf dysregulation as a key pathogenic mechanism underlying autophagy impairment in MPS VII and to demonstrate that pharmacological activation of Mitf represents a promising therapeutic strategy for restoring lysosomal-autophagy pathway and neuronal function.

## Results

### Mitf is downregulated in the CG2135^−/−^ fly brain

Autophagy defects in CG2135^−/−^ flies are evident from early adulthood which persists in following ages as examined (9). Interestingly, neurodegenerative phenotypes become pronounced at 30 and 45 days old flies as a consequential effect of the autophagy deficiency (8,9). The CG2135^−/−^ brain also showed reduced levels of autophagy markers proteins Atg8 and Ref(2)P, along with their downregulated transcripts (9). To investigate the molecular basis of autophagy deficiency, we assessed expression of Mitf, their transcriptional regulator (11), in the 30-day-old CG2135^−/−^ fly brain and compared with wildtype (WT) using RT-qPCR. Mitf transcript levels in CG2135^−/−^ brains was found to be significantly reduced by ~50% compared to WT (Fig. 1A).

**Figure 1.**
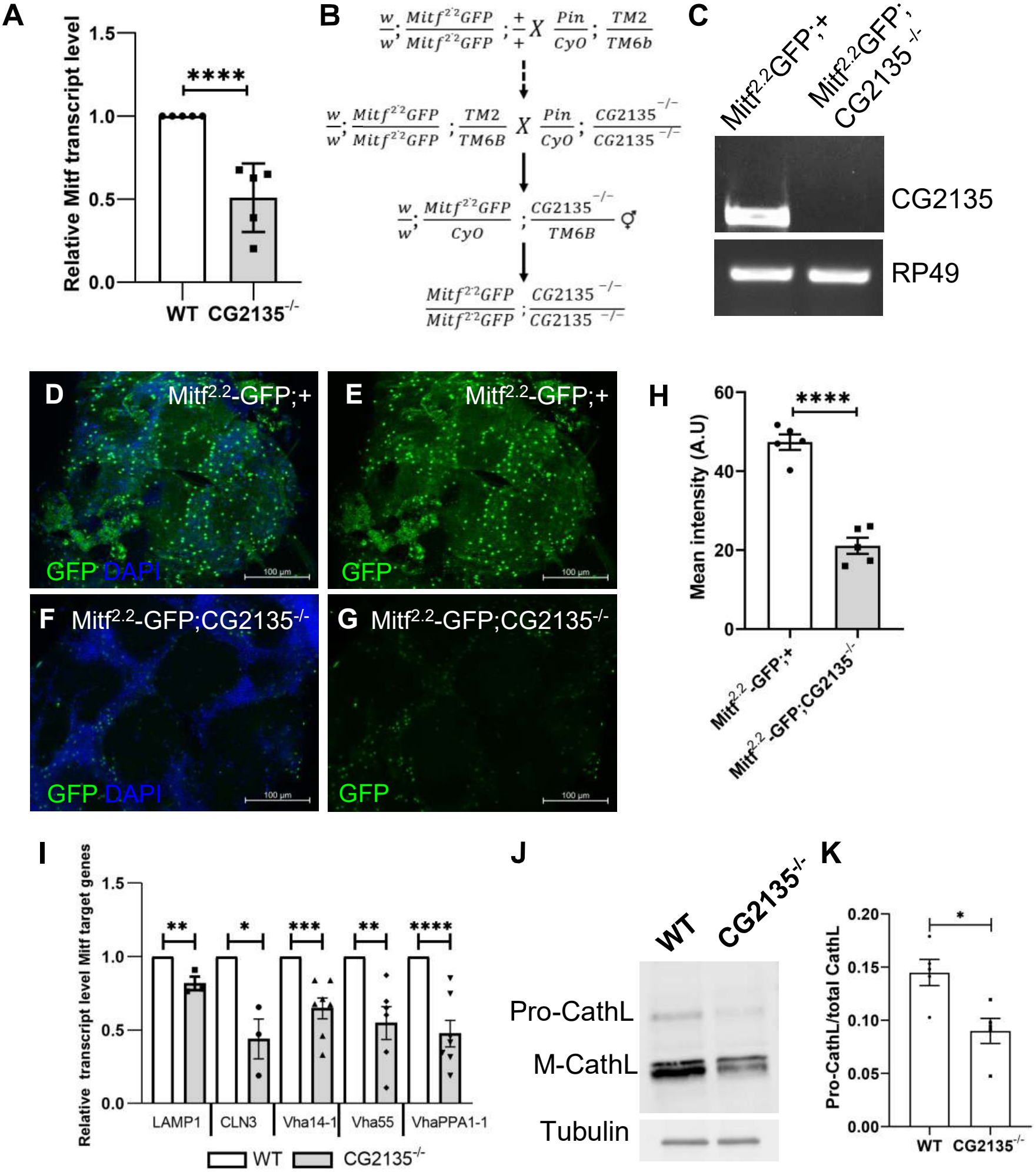
Mitf is downregulated in CG2135^−/−^ fly brain. A) Mitf transcript levels in 30-day-old WT and CG2135^−/−^ fly brain were determined by RT-qPCR and normalised to RP49 as a housekeeping gene control N = 5. B) Crossing scheme for the generation of Mitf^2·2^-GFP fly in the CG2135^−/−^ background. C) The confirmation of the Mitf^2·2^-GFP; CG2135^−/−^ fly by genomic DNA (gDNA) PCR. Agarose gel electrophoresis shows the PCR product of CG2135. The absence of the CG2135 band in Mitf^2·2^-GFP; CG2135^−/−^ fly confirms the knockout genotype, while Mitf^2·2^-GFP; + has the band. PCR with RP49-specific primers was performed as an internal control. D-G) The brain from 30-day-old Mitf^2·2^-GFP; + and Mitf^2·2^-GFP; CG2135^−/−^ shows the expression of Mitf-GFP. The green represents Mitf-GFP, and DAPI (blue) marks the nucleus. H) The bar graph represents the quantification of the GFP mean intensity, indicating the expression of Mitf N = 5 brains from each genotype. I) Transcript levels of LAMP1, CLN3, Vha14-1, Vha55, and VhaPPA1-1 in 30-day-old WT and CG2135^−/−^ fly brain were determined by RT-qPCR and normalised to RP49 as a housekeeping gene control N ≥ 3. J) Western blot of Cathepsin-L showing the protein level of pro-CathL and mature-CathL (37-25 kDa) from brain lysate of 30-day-old WT and CG2135^−/−^, as indicated in the figure. α-Tubulin (54 kDa) was used as a loading control. (K) The graph represents quantification for Pro-CathL/total CathL band intensity, showing a decreased ratio in the CG2135^−/−^ brain lysate N = 5. Error bars represent the standard error of the mean (SEM). Asterisks represent a level of significance (**** P ≤ 0.0001, *** P ≤ 0.001, ** P ≤ 0.01, * P ≤ 0.05; Student’s t-test).

To validate these finding, we used the Mitf^2·2^-GFP reporter line, in which 2.2 kb of the upstream regulatory sequence and 5′UTR of Mitf drive nuclear GFP expression (28). This promoter region contains elements responsive to endogenous Mitf activity and supports autoregulation of Mitf expression (29). To proceed with the experiment, we first generated Mitf^2·2^-GFP; CG2135^−/−^ flies through general genetic crossing scheme used in *Drosophila* model organism (Fig. 1B) and verified the knockout genotype by genomic PCR performed with CG2135-specific primers. PCR-amplification was found in WT but absence of the CG2135 PCR product confirmed the genotype of Mitf^2·2^-GFP; CG2135^−/−^ which was used for the study (Fig. 1C). Next, we examined the GFP expression in the fly digestive system, where Mitf is highly expressed (13). Fluorescence microscopy revealed GFP signals in both Mitf^2·2^-GFP; + (control) and Mitf^2·2^-GFP; CG2135^−/−^ midguts, confirming Mitf promoter activity (Fig. S1). Notably, GFP intensity was lower in Mitf^2·2^-GFP; CG2135^−/−^ midguts compared with the control. We then examined Mitf-GFP expression in the brains of 30-day-old flies (Fig. 1D-G) and observed a significant reduction in Mitf-GFP fluorescence in Mitf^2·2^-GFP; CG2135^−/−^ compared to Mitf^2·2^-GFP; + brains (Fig. 1H). Together, these results indicate a marked decrease in Mitf expression in the CG2135^−^/^−^ fly brain.

Given that Mitf regulates the expression of autophagy- and lysosome-related genes (12), its downregulation likely impacts the genes related to the CLEAR network. To test this, we quantified transcripts of selected lysosomal membrane protein genes (LAMP1, CLN3) and v-ATPase subunits Vha-14-1, Vha-55, and Vha-PPA1-1. RT-qPCR analysis of 30-day-old WT and CG2135^−/−^ brains showed significant reductions in LAMP1 and CLN3 expression, along with marked downregulation of the v-ATPase subunits (Fig. 1I). These findings indicate that Mitf downregulation is associated with reduced CLEAR network gene expression including previously reported autophagy genes Atg8a and Ref(2)P in the CG2135^−^/^−^ fly brain, highlighting Mitf’s significant role in lysosomal biogenesis and autophagy regulation in MPS VII.

Lysosomes contain hydrolases and proteases essential for degrading substrates delivered via autophagy and endocytosis. In CG2135^−^/^−^ fly brains, we observed pronounced accumulation of undegraded material, including lipofuscin and multilamellar bodies, reflecting substantial lysosomal dysfunction (9). Because Mitf regulates several v-ATPase subunits, its downregulation may alter lysosomal acidification and cause neurodegeneration (30, 31). Cathepsin-L is a cysteine protease that undergoes pH-dependent maturation from a pro-enzyme to its active form within lysosomes (32). To evaluate whether altered lysosomal acidity affects its processing, we examined pro- and mature Cathepsin-L levels in brain lysates from WT and CG2135^−/−^ flies. Immunoblot analysis revealed a decreased pro-Cath-L/total Cath-L ratio in 30-day-old CG2135^−/−^ brains relative to WT (Fig. 1J–K). This suggests that the relative conversion of pro-to mature Cathepsin-L is not impaired. However, when we analyzed the total protein levels of both pro- and mature Cathepsin-L, was found to be significantly decreased (Fig. S2A–C). This reduction indicates a loss of overall Cathepsin-L rather than a major defect in processing efficiency in the CG2135^−/−^ fly brain.

Altogether, it is evident that Mitf and its target genes are downregulated and have a role in disrupting the lysosome-autophagy pathway in the CG2135^−/−^ fly brain.

### Trehalose treatment boosts Mitf expression in the CG2135^−/−^ fly brain

We wanted to treat CG2135^−/−^ flies with a pharmacological modulator of Mitf to reverse the defect. To correct Mitf levels, we explored the potential of the small molecule trehalose, a non-reducing disaccharide of glucose have been shown to have neuroprotective properties mediated by activating TFEB (the homolog of Mitf) in mouse models of MPS II and IIIB (33, 34). Building on this evidence, we tested whether trehalose could activate Mitf levels in the CG2135^−/−^ fly brain.

Four-day-old WT and CG2135^−/−^ flies were fed standard media supplemented with 2.5% trehalose for 30–45 days, while untreated flies served as controls. We first tested the toxicity of trehalose at this dose by recording the survival of untreated WT and CG2135^−/−^ flies compared with trehalose-treated cohorts. Survival percentage plots of the flies confirmed that trehalose was non-toxic, as untreated WT and CG2135^−/−^ flies had mean lifespans of 47.7 and 38.4 days, respectively, whereas trehalose-fed WT and CG2135^−/−^ flies lived 49.6 and 38.9 days (Fig. S3A & B).

Next, to determine whether trehalose restores Mitf expression, we quantified Mitf mRNA levels in untreated and treated 30-day-old CG2135^−/−^ fly brains by RT-qPCR and compared them with untreated WT fly brain. Trehalose treatment completely rescued Mitf transcript levels in the CG2135^−/−^ fly brain similar to WT levels (Fig. 2A). We next validated the trehalose-mediated Mitf restoration using the Mitf^2.2^-GFP reporter line, which reports endogenous Mitf transcriptional activity. Four-day-old Mitf^2.2^-GFP; CG2135^−/−^ flies were fed trehalose-supplemented media, while untreated Mitf^2.2^-GFP; + and Mitf^2.2^-GFP; CG2135^−/−^ flies served as controls. After 30 days, fluorescence imaging revealed a marked increase in the Mitf-GFP signal in trehalose-treated Mitf^2.2^-GFP; CG2135^−/−^ brains compared to untreated counterparts, confirming the restoration of Mitf activity (Fig. 2B–G). Figure 2H shows the quantification of the GFP intensity in untreated Mitf^2.2^-GFP; + and Mitf^2.2^-GFP; CG2135^−/−^ and trehalose-treated Mitf^2.2^-GFP; CG2135^−/−^.

**Figure 2.**
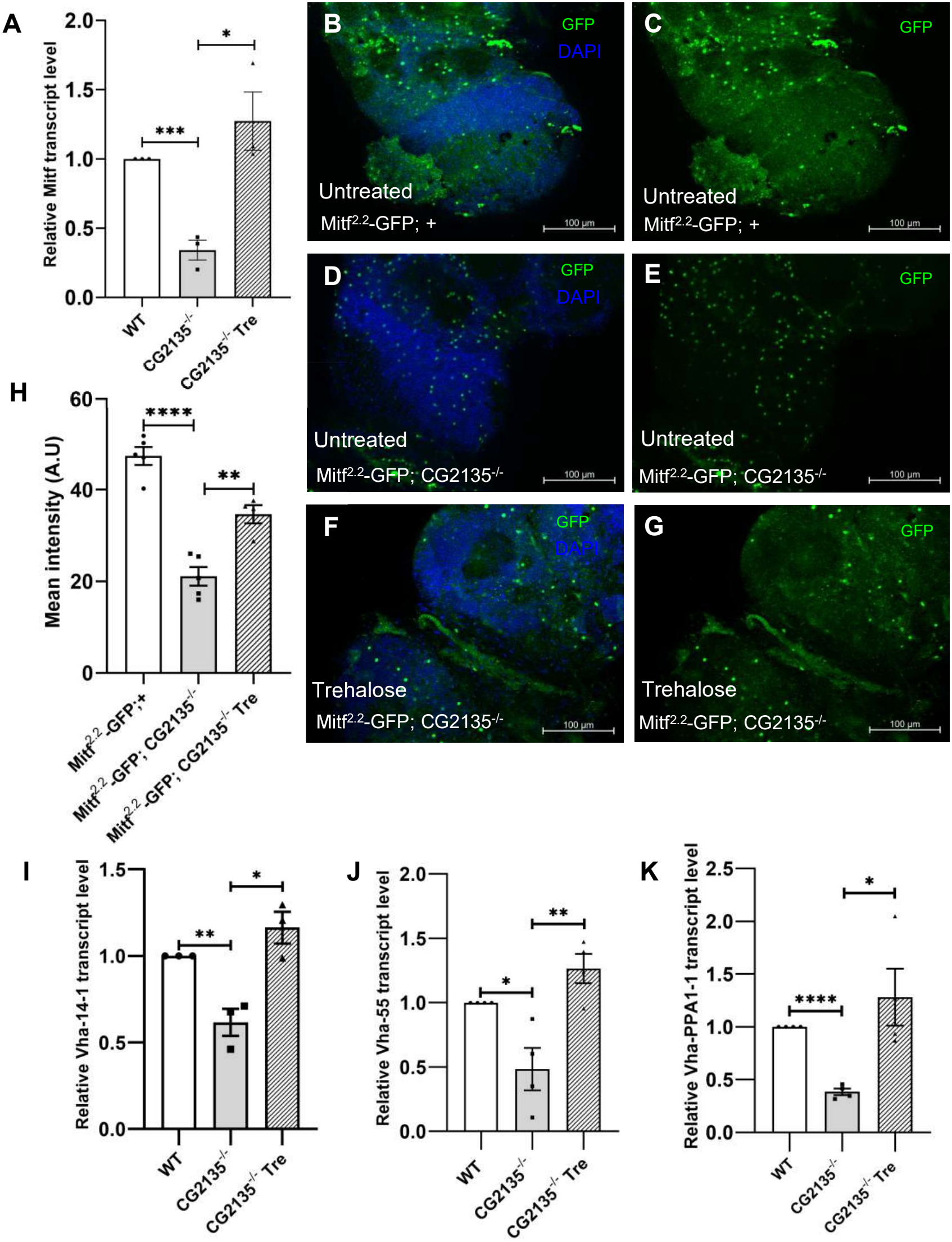
Trehalose induces Mitf expression in CG2135^−/−^ fly brain. A) Mitf transcript levels in 30-day-old untreated WT, CG2135^−/−^, and trehalose-treated CG2135^−/−^ fly brain were determined by RT-qPCR and normalised to RP49 as a housekeeping gene control N = 3. B-G) The brain from 30-day-old Mitf^2·2^-GFP; +, Mitf^2·2^-GFP; CG2135^−/−^, and trehalose-treated Mitf^2·2^-GFP; CG2135^−/−^ shows the expression of Mitf-GFP, where green puncta represent Mitf-GFP, and blue marks mark the nucleus stained with DAPI. H) The bar graph represents the quantification of the GFP mean intensity, indicating the expression of Mitf N = 5 brains from each Mitf^2·2^-GFP; +, Mitf^2·2^-GFP; CG2135^−/−^, and trehalose-treated Mitf^2·2^-GFP; CG2135^−/−^). I) Vha14-1 transcript levels in 30-day-old untreated WT, CG2135^−/−^, and trehalose-treated CG2135^−/−^ fly brain were determined by RT-qPCR and normalised to RP49 as a housekeeping gene control N = 3. J) Vha-55 transcript levels in 30-day-old untreated WT, CG2135^−/−^, and trehalose-treated CG2135^−/−^ fly brain were determined by RT-qPCR and normalised to RP49 as a housekeeping gene control. K) Vha-PPA1-1 transcript levels in 30-day-old untreated WT, CG2135^−/−^, and trehalose-treated CG2135^−/−^ fly brain were determined by RT-qPCR and normalised to RP49 as a housekeeping gene control. Error bars represent the standard error of the mean (SEM). Asterisks represent a level of significance (**** P ≤ 0.0001, *** P ≤ 0.001, ** P ≤ 0.01, * P ≤ 0.05; Student’s t-test).

Next, we examined the expression same of the Mitf target genes as v-ATPase subunits Vha-14-1, Vha-55, and Vha-PPA1-1. RT-qPCR analysis revealed that the transcript levels of these genes were restored to normalcy us, untreated WT levels, indicating upregulation of all three genes in trehalose-treated CG2135^−/−^ brains relative to untreated CG2135^−/−^ fly brains (Fig. 2I–K).

Collectively, these findings demonstrate that trehalose treatment reinstates Mitf expression and activity in CG2135^−/−^ flies, thereby rescuing the transcription of downstream lysosomal genes. trehalose thus can be used as a therapeutic to restore Mitf-mediated autophagy-lysosomal function in MPS VII.

### Trehalose activates autophagy and restores mTOR signaling in the CG2135^−/−^ fly brain

We previously demonstrated that defective autophagy drives neurodegeneration in CG2135^−/−^ flies (9). Given that Mitf dysregulation underlies this defect, we next examined whether trehalose could restore autophagy via Mitf activation. Activation of autophagy initiates by the formation of a vesicle known as the autophagosome, and its maturation requires the Mitf target gene Atg8a. Atg8a is a cytosolic protein which undergoes post-translational modification. The membrane-bound form, Atg8a-II, is processed from Atg8a-I by other Atg proteins, such as Atg4, and recruited to the autophagosomes (35). An increase in the Atg8a-II levels reflects autophagy activation and autophagosome formation, while a decrease in the Atg8a-II may indicate its autophagic degradation in the lysosomes (36). Based on this rational, we assessed autophagy induction and measured Atg8a-II protein levels—a marker of autophagosome formation—in brain lysates from 30-day-old untreated and trehalose-fed WT and CG2135^−/−^ flies using immunoblotting.

In WT flies, trehalose treatment decreased Atg8a-II levels compared to untreated controls (Fig. 3A–B), consistent with enhanced autophagic flux and increased autophagosome turnover by lysosomes. In contrast, CG2135^−/−^ flies displayed a marked increase in Atg8a-II upon trehalose treatment relative to untreated mutants, indicating autophagy induction in the diseased background. These results suggest that trehalose stimulates autophagy in both WT and CG2135^−/−^ brains. The upregulation of Atg8a levels was corroborated by increase in the transcript levels upon trehalose treatment, as determined by RT-qPCR comparing untreated and trehalose-treated CG2135^−/−^ fly brains (Fig. 3C). Additionally, the Mitf target gene Ref(2)P was also upregulated in trehalose-treated CG2135^−/−^ fly brains (Fig. 3D). Moreover, the restored levels of the autophagy genes and proteins are like the untreated WT flies. Next, we determined the molecular basis of autophagy induction by assessing the transcript levels of *Atg1*, a key autophagy-initiating kinase (37). *Atg1* expression was significantly upregulated in trehalose-treated CG2135^−/−^ brains compared with untreated mutants (Fig. 3E), suggesting that trehalose triggers autophagy initiation through *Atg1* activation. Together, these findings demonstrate that trehalose enhances autophagy markers by upregulating *Atg1* and Mitf-dependent autophagy genes (*Atg8a* and *Ref(2)P*), and is consistent with increased autophagosome formation in CG2135^−/−^ flies.

**Figure 3.**
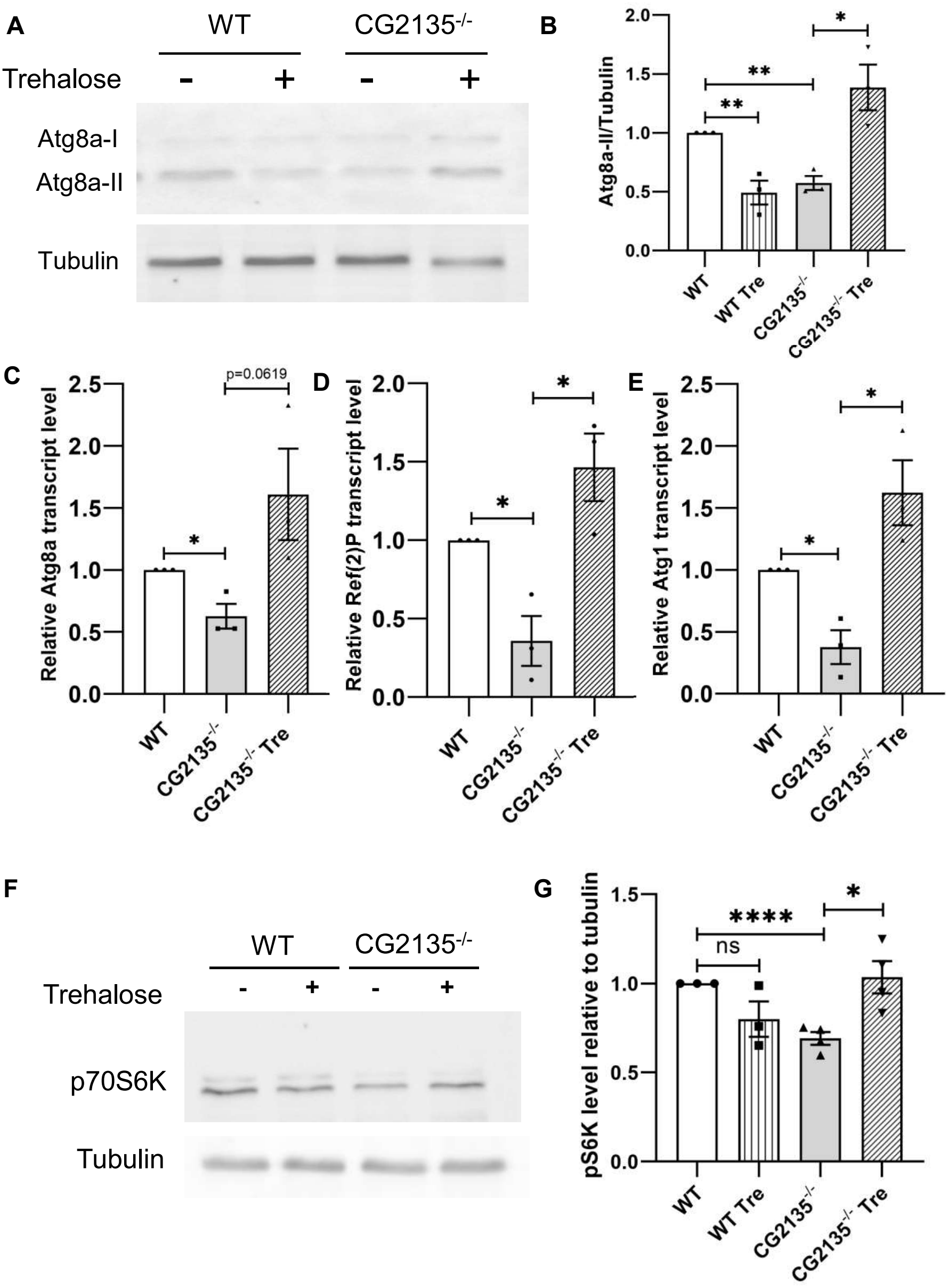
Trehalose restores the autophagy defect in the CG2135^−/−^ fly brain. A) Western blot of Atg8a showing the protein level of Atg8a-II (14 kDa) from brain lysate of 30-day-old untreated WT and CG2135^−/−^ and trehalose-treated WT and CG2135^−/−^ fly brains. (B) The bar graph represents quantification for Atg8a-II/Tubulin band intensity, showing an increased level of Atg8a-II in the trehalose-treated CG2135^−/−^ head lysate compared to untreated CG2135^−/−^ N = 3. C) Atg1 transcript levels in 30-day-old untreated WT, CG2135^−/−^, and trehalose-treated CG2135^−/−^ fly brain were determined by RT-qPCR and normalised to RP49 as a housekeeping gene control N = 3. D) Atg8a transcript levels in 30-day-old untreated WT, CG2135^−/−^, and trehalose-treated CG2135^−/−^ fly brain were determined by RT-qPCR and normalised to RP49 as a housekeeping gene control N = 3. E) Ref(2)P transcript levels in 30-day-old untreated WT, CG2135^−/−^, and trehalose-treated CG2135^−/−^ fly brain were determined by RT-qPCR and normalised to RP49 as a housekeeping gene control N = 3. F) Scheme for the canonical pathway of mTOR signalling and experimental set-up to study the mTOR level. B) Western blot of p70S6K showing the protein level kDa) from brain lysate of 30-day-old untreated WT and CG2135^−/−^ and trehalose-treated WT and CG2135^−/−^ fly brains. α-Tubulin (54 kDa) was used as a loading control. (G) The graph shows quantification of p70S6K/Tubulin band intensity, indicating restored mTOR levels in the trehalose-treated CG2135^−/−^ head lysate N = 3. Error bars represent the standard error of the mean (SEM). Asterisks represent a level of significance (**** P ≤ 0.0001, ** P ≤ 0.01, * P ≤ 0.05; Student’s t-test).

We next examined whether trehalose-induced autophagy depends on mTOR regulation. mTOR is a central metabolic regulator that integrates nutrient sensing with proteostasis and lysosomal function, and it tightly controls both autophagy and the activity of the transcription factor Mitf (38). Accordingly, mTOR activity in untreated and trehalose-treated CG2135^−/−^ fly brains and corresponding WT controls was assessed by measuring the phosphorylation of its downstream substrate, p70S6 kinase (p70S6K), at Thr398 (39). Immunoblot analysis revealed reduced phosphorylated p70S6K (p-p70S6K) levels in untreated CG2135^−/−^ brains compared with untreated WT brains, indicating decreased mTOR activity (Fig. 3F–G). Interestingly, trehalose treatment significantly increased p70S6K levels in the CG2135^−/−^ fly brains relative to their untreated counterparts, restoring mTOR activity toward near-WT levels. This restoration of mTOR signalling occurred alongside the recovery of Mitf/TFEB expression and autophagy described above, suggesting that trehalose improves cellular homeostasis in the CG2135^−/−^ brain by re-establishing both lysosomal-autophagy pathways and mTOR-dependent metabolic signalling.

### Trehalose mitigated neurodegeneration in CG2135−/^−^ flies and restored their locomotor activity

The brain, composed primarily of post-mitotic neurons, relies heavily on the autophagy-lysosomal pathway for the clearance of age-related aggregates such as lipofuscins (40, 41). We have demonstrated that trehalose restores Mitf levels-the master regulator of autophagy and lysosomal gene networks—in CG2135^−/−^ brains. Since Mitf activation is known to enhance lysosomal biogenesis and exocytosis (42, 43), we hypothesised that trehalose-induced Mitf activation would facilitate the clearance of lipofuscin-like autofluorescent aggregates in CG2135^−/−^ brains.

To test this, brains from 45-day-old untreated WT, CG2135^−/−^, and trehalose-treated CG2135^−/−^ flies were dissected and imaged under the FITC channel to visualise autofluorescent bodies. All images were acquired at identical settings and quantified using a defined threshold in ImageJ. Interestingly, trehalose-treated CG2135^−/−^ brains exhibited a significant reduction in autofluorescent aggregates compared with untreated CG2135^−/−^ brains (Fig. 4A–D), indicating enhanced lysosomal clearance activity.

**Figure 4.**
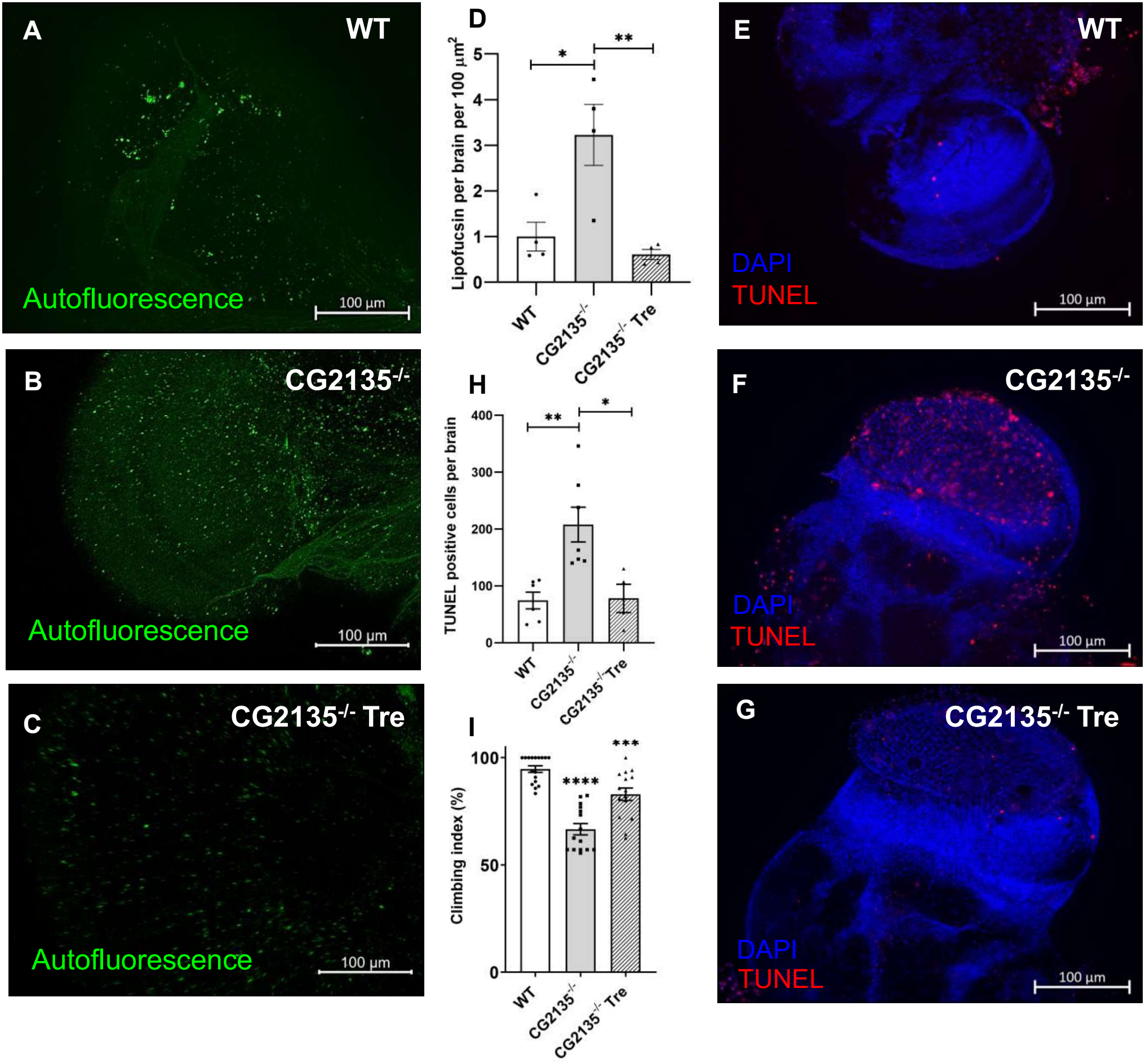
Trehalose corrects the neurodegeneration in the CG2135^−/−^ fly brain. A-C) Autofluorescent bodies known as lipofuscins (detected using the FITC channel) in 45-day-old untreated WT and CG2135^−/−^ and trehalose-treated CG2135^−/−^ fly brains, showing an enhanced clearance of autofluorescent bodies in trehalose-treated CG2135^−/−^. D) Bar graph depicting mean autofluorescence intensity of WT, CG2135^−/−^, and trehalose-treated CG2135^−/−^ fly brains N = 4. E-G) TUNEL staining (red) of 45-day-old WT, CG2135^−/−^, and trehalose-treated CG2135^−/−^ fly brains, with DAPI (blue) used for marking the nucleus. (H) The bar graph shows quantification of the TUNEL-positive nuclei, showing a decrease in TUNEL-positive cells in the trehalose-treated CG2135^−/−^ fly brain indicating restoration of neuronal health N ≥ 4 fly brains). I) Climbing index of 30-day-old untreated WT, CG2135^−/−^, and trehalose-treated CG2135^−/−^ fly brains showing improved climbing ability in trehalose-treated CG2135^−/−^ flies N = 120 flies. Error bars represent the standard error of the mean (SEM). Asterisks represent a level of significance (**** P ≤ 0.0001, *** P ≤ 0.001, ** P ≤ 0.01, * P ≤ 0.05; Student’s t-test).

Next, we evaluated whether Mitf activation and improved lysosomal function were associated with reduced apoptosis, the underlying cause of neurodegeneration in MPS VII (9). Whole-mount TUNEL staining was performed on brains from 45-day-old untreated WT, CG2135^−/−^, and trehalose-treated CG2135^−/−^ flies. Trehalose-treated CG2135^−/−^ brains showed a marked reduction in TUNEL-positive cells compared with untreated CG2135^−/−^ fly brains (Fig. 4E-H), signifying decreased neuronal apoptosis.

Finally, to assess whether trehalose-mediated molecular restoration translates to functional improvement, we evaluated the climbing index, a behavioural marker of neurodegeneration reported in CG2135^−/−^ flies (Bar et al., 2018). The 30-day-old trehalose-treated CG2135^−/−^ flies exhibited a significantly improved climbing ability compared to untreated mutants (Fig. 4I).

Together, these results demonstrate that trehalose rescues lysosomal-autophagy function, reduces apoptosis, and improves locomotor performance in CG2135^−/−^ flies, underscoring its therapeutic potential in alleviating neurodegeneration associated with MPS VII.

## Discussion

Correction of neuropathology in MPS VII remains an unmet therapeutic challenge. A limited understanding of the molecular mechanisms underlying neurodegeneration in MPS VII has hindered the development of CNS-directed therapy. In this study, we identify Mitf suppression as a key molecular defect in the MPS VII brain and demonstrate that pharmacological activation of Mitf by trehalose restores autophagy, ameliorates neurodegenerative phenotypes and restores locomotor function in the CG2135^−/−^ flies (summarized in Fig. 5). Collectively, these findings establish Mitf activation as a safe and targetable therapeutic mechanism capable of correcting neuronal pathology, which current enzyme replacement therapy fails to achieve.

TFEB/Mitf plays a crucial role in regulating autophagy-lysosome biogenesis to maintain neuronal health (11,12). Reduced expression of TFEB has been reported in monocytes and lymphocytes from Alzheimer’s diseases (AD) patients, as well as in mouse models of Amyotrophic lateral sclerosis (ALS), and Huntington’s disease (HD) (20,44). In Gaucher disease iPSC derived neuron cells, significant downregulation of TFEB was found while its *Drosophila* model showed upregulation in brain tissues (45,46). Both conditions led to impaired autophagy associated with neurodegeneration demonstrating a direct correlation between expression of CLEAR network linked autophagy-lysosome related genes and TFEB/Mitf levels. Impairment in the TFEB-v-ATPase regulatory axis has been linked to abnormal tau accumulation and defective microglial activation in AD (47). Similarly, loss of the v-ATPase subunit ATP6AP2 induces neurodegeneration and cognitive decline in both fly and mouse models of Parkinson’s disease (PD), accompanied accumulation by autophagic vacuoles (48,49). Altered v-ATPase levels disrupt acidification of lysosomes, thereby impairing lysosomal hydrolases and proteases (50). The CG2135^−/−^ fly brain, no apparent defect in cathepsin-L processing was observed, in contrast to the *Drosophila* ALS model in which reduced TFEB activity was shown to impair cathepsin-L maturation (51). However, reduced cathepsin-L expression in the CG2135^−/ −^ fly brains may arise from compromised lysosomal membrane stability and broader dysregulation of lysosomal protein expression, as evidenced by neuronal vulnerability reported in the lysosomal storage disease Niemann-Pick disease type C1 (52,53). Supporting our findings, the accumulation of multilamellar bodies and lipofuscin (9) reinforces that lysosomal degradative capacity is compromised, leading to buildup of undegraded materials and defective autophagic turnover. Recent studies have provided evidence that post-transcriptional modifications, including miRNA-mediated regulation, as well as acetylation, and deacetylation to influence activity and localisation of TFEB/Mitf (54). Interestingly, downregulation of TFEB/Mitf through acetylation by histone acetyltransferase GCN5 supress its transcriptional activity leading to autophagy impairment in Hela cells and in hemocytes isolated from *Drosophila* (55,56). In future, investigating post-transcriptional regulatory mechanisms in the context of MPS VII will advance our understanding of how lysosomal dysfunction interferes with downstream cellular signalling.

**Figure 5.**
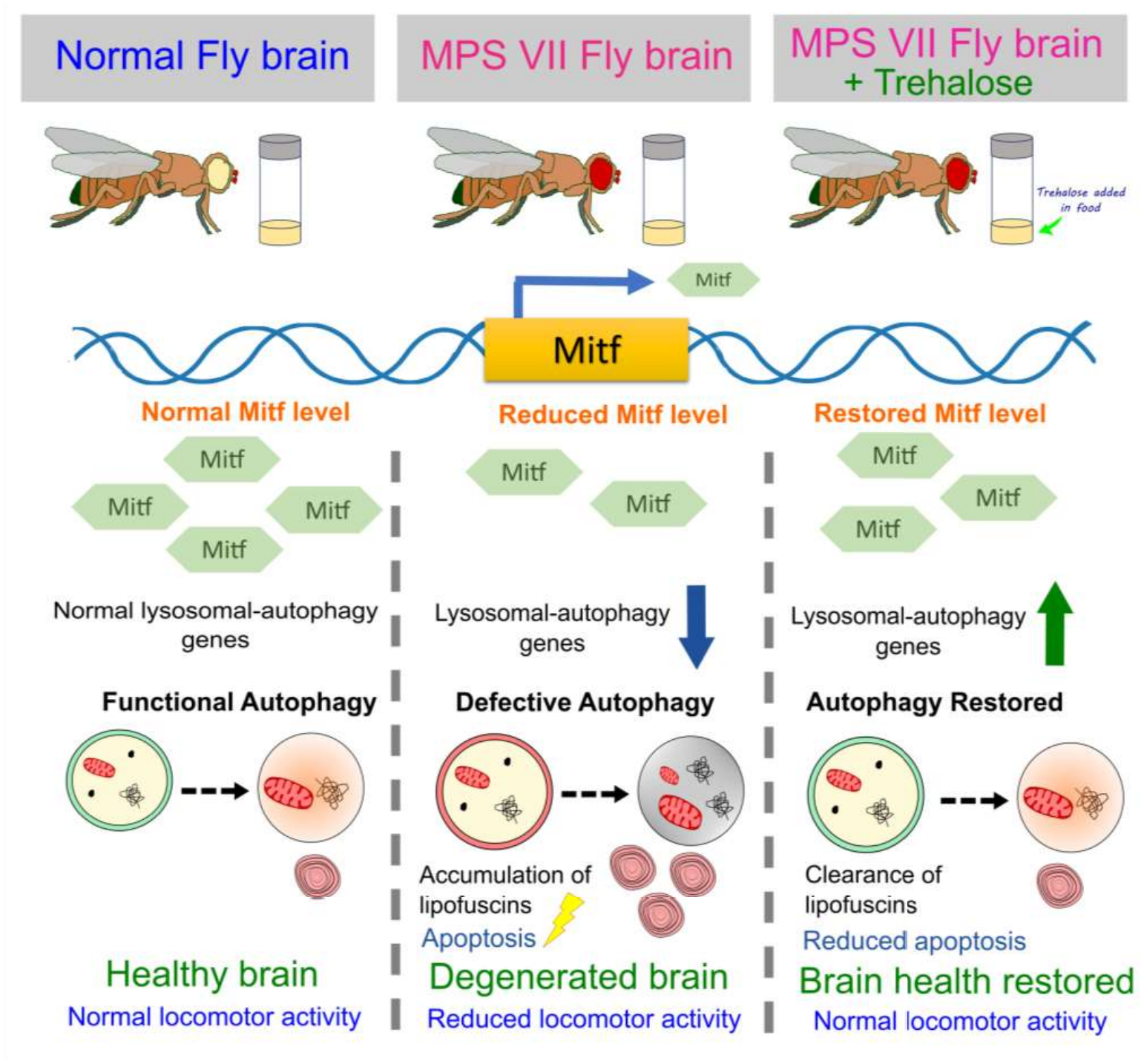
Trehalose-mediated restoration of Mitf corrects autophagy defects and ameliorates neurodegeneration in MPS VII flies. In the normal fly brain (left), normal levels of the transcriptional regulator Mitf maintain the expression of lysosomal-autophagy genes, ensuring functional autophagy and normal locomotor activity. In the MPS VII fly brain (centre), Mitf levels are markedly reduced, leading to a down-regulation of the autophagy gene network. This results in defective autophagy, characterized by the toxic accumulation of lipofuscins and damaged mitochondria, which triggers apoptosis, brain degeneration, and impaired locomotor performance. Pharmacological intervention via oral trehalose treatment (right) restores Mitf levels and reactivates the downstream autophagic machinery in the MPS VII flies. This systemic recovery enhances cellular clearance, reduces apoptotic cell death, and restores brain health and locomotor activity of the MPS VII flies to near-normal levels.

In untreated CG2135^−/−^ fly brains, mTOR activity is markedly reduced, accompanied by decreased Mitf expression and downregulation of its target genes, including Atg8, Ref(2)P/p62, and v-ATPase subunits resulting in impaired autophagic flux and progressive neurodegeneration. Trehalose treatment restored transcriptional levels of Mitf and its target autophagy genes Atg8 and Ref(2)P, occurring concomitantly with increased mTOR activity in CG2135^−/−^ fly brain. This observation challenges the canonical model in which mTOR activation suppresses TFEB/Mitf and autophagy (57–59). Interestingly, a similar paradoxical coexistence of active TFEB and elevated mTOR signaling has been reported in diseases such as pancreatic cancer and Birt-Hogg-Dubé syndrome, where coordinated activation of autophagy and lysosomal catabolism supports metabolic adaptation and cell survival under nutritional stress (60–64). In MPS VII, lysosomal dysfunction likely creates a state of metabolic insufficiency in which both TFEB and mTOR signaling are suppressed. Trehalose treatment may restore lysosomal competence through TFEB activation while simultaneously improving metabolic capacity via mTOR signaling. Thus, in the CG2135^−/−^ brain, increased mTOR activity during recovery does not inhibit TFEB driven autophagy but instead reflects improved cellular metabolic fitness. Collectively, these findings broaden the understanding of lysosomal mTOR crosstalk in neurodegeneration and highlight TFEB activation as a potential therapeutic strategy to restore autophagic function in MPS VII.

The potential of trehalose in activating TFEB to correct neurodegeneration has been shown in pre-clinical models of Huntington’s disease, dementia, Batten disease, MPS IIIB and recently in MPS II among others (34,65–68). The study in MPS II lacked molecular mechanism while MPS IIIB showed trehalose rescued retinal degeneration through TFEB activation in treated mutant mouse but there is no difference of basal TFEB expression in untreated mutant in comparison to wildtype mouse. Thus, our work reinforces the importance of systemic investigation of Mitf and its modulation using trehalose to alleviate neuropathology in MPS VII brain. TFEB/Mitf being a transcriptional factor its activity depends on cytosol and nucleus localisation (54) where trehalose has the potential to modulate TFEB/Mitf activity which promotes cellular debris clearance through lysosomal exocytosis or enhanced lysosomal activity (69). Mechanistically, trehalose has been shown to induce TFEB/Mitf nuclear localisation, which is essential for its transcriptional activity by promoting dephosphorylation that disrupts its interaction with 14-3-3 protein by phosphatase calcineurin due to release of Ca^2+^ from the lysosomes (26), and inhibiting Akt-mediated phosphorylation of TFEB (Palmieri et al., 2017). In addition, trehalose activates autophagy through m-TOR independent pathways by upregulating autophagy-lysosome related genes, demonstrated in amelioration of signs of neurodegeneration associated with alpha-synuclein accumulation as well as in Batten disease and ALS (67,68,70). Collectively, based on accumulating evidence of its beneficial effects and established safety and tolerability in patients, trehalose is currently being evaluated in clinical trials at various stages for diseases like spinocerebellar ataxia type 2 (stage II/III), Parkinson’s (stage IV), Alzheimer’s (stage II), ALS (II/III) etc (71,72). Notably, trehalose can also cross the blood brain barrier, enable direct central nervous system exposure, and contribute to the attenuation of neuropathology. Thus, our findings support trehalose administration as a promising therapeutic strategy to correct neurodegeneration in MPS VII.

In conclusion, our study links lysosomal storage pathology to impaired transcriptional control of cellular clearance pathways, providing a mechanistic and therapeutically tractable framework for neuronopathic MPS VII. Given trehalose’s clinical safety and translational progress, these findings support Mitf/TFEB modulation as a promising strategy for CNS-targeted intervention in lysosomal storage diseases.

## Material and methods

All reagents were procured from Sigma Aldrich unless otherwise mentioned and oligonucleotides from IDT

### *Drosophila* strains and maintenance

Wild-type W^1118^ (WT) flies were obtained from the Bloomington *Drosophila* Stock Center (BDSC), Indiana University, USA, and the CG2135^−/−^ mutant line was generated and characterised previously in our laboratory (8). Mitf^2·2^GFP; + line was gifted by Prof. Francesca Pignoni, SUNY Upstate Medical University. Flies were maintained under standard laboratory conditions at 25°C with a 12-hour light/dark cycle and in controlled density. Flies were reared on standard cornmeal medium composed of corn flour (75 g/L), sugar (80 g/L), agar (10 g/L), dry yeast (15 g/L), and malt extract (30 g/L). To preserve genetic consistency, fly stocks were regularly outcrossed before experimental use.

For obtaining age-matched flies, adults were allowed to lay eggs for 24 hours in synchronized cage setups, after which newly hatched larvae were transferred in equal numbers into food vials. Following 9-10 days of development, newly eclosed adult flies were collected and used for experiments. Only male flies were selected for experiments to minimize sex-dependent variability.

For drug treatment, compounds were mixed into cool, molten standard media before solidification. Trehalose (cat. no. 79449, Sigma) was added to the food at a final concentration of 2.5%. Treatment began on 4-day-old flies and was administered every alternate day for a period of 30 or 45 days. Freshly prepared drug-containing media were used for each feeding to ensure uniform exposure. Flies were maintained in groups of 10-20 males per vial, with each strain divided into treated and untreated cohorts for comparison.

### Genomic DNA isolation

Approximately 5-10 flies were homogenized in 200 µL of Buffer A (100 mM Tris-Cl (pH 7.5), 100 mM EDTA, 100 mM NaCl, and 0.5% SDS) using a micro-pestle. After homogenization, an additional 200 µL of Buffer A was added, and the lysate was incubated at 65°C for 30 minutes. Subsequently, 800 µL of Buffer B (5M Potassium acetate and 6M lithium chloride) was added, and the mixture was incubated at room temperature (RT) for 10 minutes. The sample was then centrifuged at 12,000 rpm for 15 minutes at RT, and the resulting supernatant (1 mL) was carefully transferred to a fresh 2 mL microcentrifuge tube.

An equal volume of phenol:chloroform (1:1) was added, mixed thoroughly, and the solution was incubated at RT for 10 minutes, followed by centrifugation at 12,000 rpm for 15 minutes at 4°C. The aqueous phase was collected into a new tube, to which 600 μL of isopropanol was added. The mixture was gently inverted to mix and centrifuged at 12,000 rpm for 15 minutes at RT to pellet the DNA. The pellet was then washed with 1 mL of 70% ethanol, air-dried after ethanol removal, and finally resuspended in nuclease-free water.

DNA quality and concentration were assessed using a NanoDrop 2000 spectrophotometer, and samples were stored at −20°C until further use.

### *Drosophila* climbing assay

On the day prior to the experiment, flies were sorted into separate batches of 20 individuals each and maintained in fresh food vials under standard culture conditions for 24 hours. On the day of the assay, flies were gently transferred to a graduated 50 mL measuring cylinder without anaesthesia. The mouth of the cylinder was sealed with a cotton plug, and the 15 mL mark was clearly indicated using a marker.

For the negative geotaxis (climbing) assay, a minimum of 10 batches of 20 flies each was tested. After a 2-minute acclimation period, flies were tapped down to the bottom of the cylinder, and the number of flies that climbed above the 15 mL mark within 15 seconds was recorded. The climbing index was calculated as the percentage of flies that successfully reached the 15 mL mark within the specified time.

The experiment was performed independently several times, and the results were represented as combined data from all biological replicates.

### Lifespan assay

A total of 200 healthy male flies were collected on the day of eclosion and transferred into six standardised culture vials. Flies were maintained for three days under normal growth conditions to allow natural reproductive activity. After this period, flies were separated into new vials, with no more than 20 flies per vial. The vials were changed every alternate day, and the number of dead flies was recorded at each transfer. Monitoring continued until all flies had died. The percentage of survivability was determined based on the ratio of live flies to the total number of flies at each time point. Mean lifespan was measured following the method described by (73).

### Western blot

For western blot, 5-10 fly whole brains were homogenized in 1× Laemmli’s (for 1 ml 2 × 0.5 M Tris-Cl pH 6.8 0.312 ml, SDS 0.05 g, glycerol 0.25 ml, 1 M DTT 50 μl) buffer supplemented with 1× protease inhibitor cocktail and boiled at 100°C for 5 min. An equal volume of the brain lysate was subjected to SDS-PAGE. Gels were transferred for 1 h and blocked in 5 % skim milk in 0.025 % TBST. The primary antibody (anti-GABARAP cat no. E1J4E, CST; anti-Cathepsin-L cat no. MAB22591, RnD systems; anti–p70S6K cat no. p1705–398, PhosphoSolutions; anti-α-Tubulin cat no. AB0118, BioBharati) incubation was done at 4 °C overnight in respective dilutions of antibodies. The secondary antibodies (Goat Anti-Rabbit IgG-Peroxidase, cat no. A9169, Sigma; Goat Anti-Mouse IgG-Peroxidase, cat no. A9044, Sigma) were incubated after washing for 1.5 h. After final washing, the blots were developed using Super Signal West Pico Chemiluminescent Substrate (Thermo Fisher) and captured using the ChemiDoc imaging system (Biorad). α-Tubulin has been used as a loading control. Densitometric analysis of protein bands was quantified using ImageJ software.

### RNA isolation and RT-qPCR

Total RNA was extracted from 10-20 fly whole brains of WT and CG2135^−/−^ strains using TRIzol reagent (Invitrogen), following the manufacturer’s protocol. A total of 1 μg of RNA was treated with DNase to remove residual genomic DNA, and the purified RNA was subsequently used for cDNA synthesis using a first-strand cDNA synthesis kit. The resulting cDNA served as the template for quantification of gene transcripts. Quantitative real-time PCR (qRT-PCR) was carried out using SYBR Green PCR Master Mix (Bio-Rad) on a Bio-Rad real-time PCR detection system. Expression levels of target genes (primer sequences listed in Table S1) were normalized to the reference gene RP49 as an endogenous control. Relative gene expression was calculated using the comparative threshold cycle (ΔΔCT) method (74).

### TUNEL staining

To assess cell death in WT and CG2135^−/−^ fly brains, a TUNEL assay (cat no. 12156792910, Roche) was performed as described by (75). Dissected brains were collected in PBS and fixed in 4% paraformaldehyde (PFA) containing 0.1% Triton X-100 for 30 minutes. Samples were then washed twice each in 0.1% PT and 0.5% PT for 3 minutes per wash. The brains were permeabilized in sodium citrate at 65°C for 30 minutes, followed by three washes in 0.5% PT. The tissues were then incubated with the TUNEL reaction mix at 37°C for 3 hours, following the manufacturer’s protocol. After incubation, samples were washed three times with 0.1% PT, counterstained with DAPI, and mounted in Vectashield for imaging. Images were acquired the following day using a Zeiss Apotome.2 fluorescence microscope and processed with ZEN software. For each brain, 15-20 z-stacks were captured to generate maximum intensity projections (MIPs). TUNEL-positive puncta were quantified after thresholding in ImageJ, and the count was expressed per brain.

### Lipofuscin imaging and analysis

For lipofuscin imaging, the procedure was adapted from (76). Age-matched fly brains were dissected in cold phosphate buffer and kept on ice for 20 minutes. The samples were then immediately imaged using a Zeiss Apotome.2 microscope under the FITC channel to capture autofluorescence signals. Images were processed using ZEN Blue software. For quantification, maximum intensity projections (MIPs) were generated from 15 z-stacks, and a uniform threshold was applied across all samples. The number of autofluorescent (lipofuscin) bodies was determined using ImageJ analysis software.

### Mitf^2.2^GFP fly tissue dissection, image acquisition and analysis

Reporter line Mitf^2.2^GFP flies were reared in standard fly food. On the day of experiment, tissue as midgut and brain was dissected from the flies in S2 media supplemented with 10 % FBS (Avantor), 100 U/ml penicillin (Gibco) and 100 μg/ml streptomycin (Gibco). After tissue isolation, they were fixed in 4% paraformaldehyde (PFA) for atleast 20 minutes. Then they were washed with PBS and transferred in 0.3% PBT (PBS containing 0.3% triton-X 100). The tissues were stained in DAPI (1 μg/ml) for 20 mins and mounted in Vectashield and stored at 4°C. Images were acquired in Apotome.2 microscope and images processed in Zen blue from Carl Zeiss. 15-20 Z planes were merged as MIP (maximum intensity projection) and the mean was taken from the multiple ROIs for each field and quantified in ImageJ after applying same threshold to determine the mean intensity of the GFP of the tissues.

### Statistical analysis

All the experiments were done at least three times independently. All the graphs have been plotted and statistically analysed using GraphPad Prism. *p*-values ≤0.05 were considered statistically significant as indicated by asterisks: *p≤0.05, p≤0.01, ***p≤0.001 and ****p≤0.0001.

## Acknowledgements

The authors express their sincere gratitude to Prof. Mohit Prasad and the members of fly-lab for providing fly facility. Abhrajyoti Nandi, Sujoy Bose and Subhajit Majumdar are appreciated for their technical assistance.

## Author contributions

Apurba Das: Writing - Original Draft, Methodology, Investigation, Formal Analysis, Data Curation, Conceptualization. Rupak Datta: Writing - Original Draft, Review & Editing, Supervision, Resources, Project Administration, Funding Acquisition, Formal Analysis, Data Curation, Conceptualization.

## Data Availability Statement

All data are fully described in the manuscript and supplementary materials

## Supplementary Information

**Figure S1:**
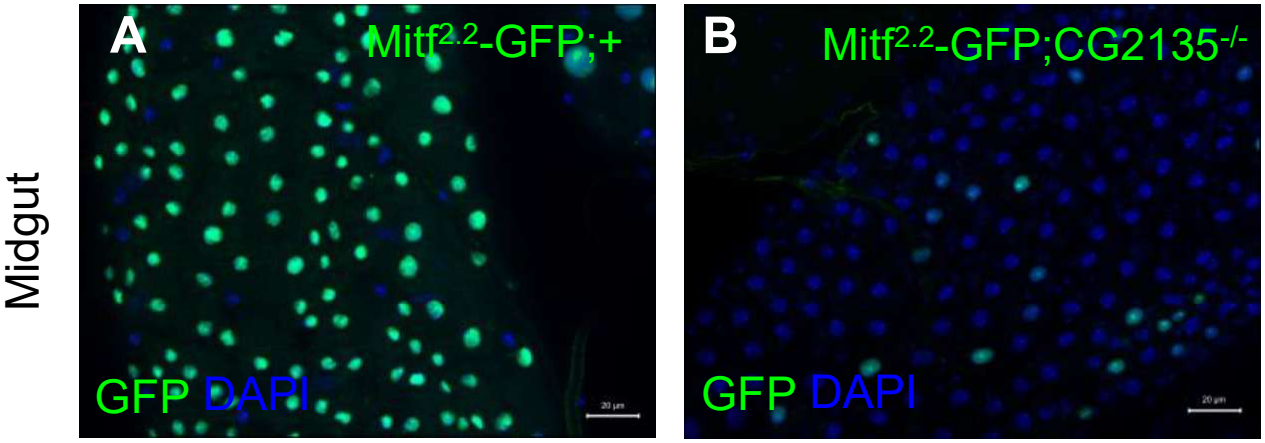
(A-B) The midgut from Mitf^2·2^-GFP; + and Mitf^2·2^-GFP; CG2135^−/−^ shows the expression of Mitf-GFP, validating the reporter line expression. The green represents Mitf-GFP, and DAPI (blue) marks the nucleus.

**Figure S2:**
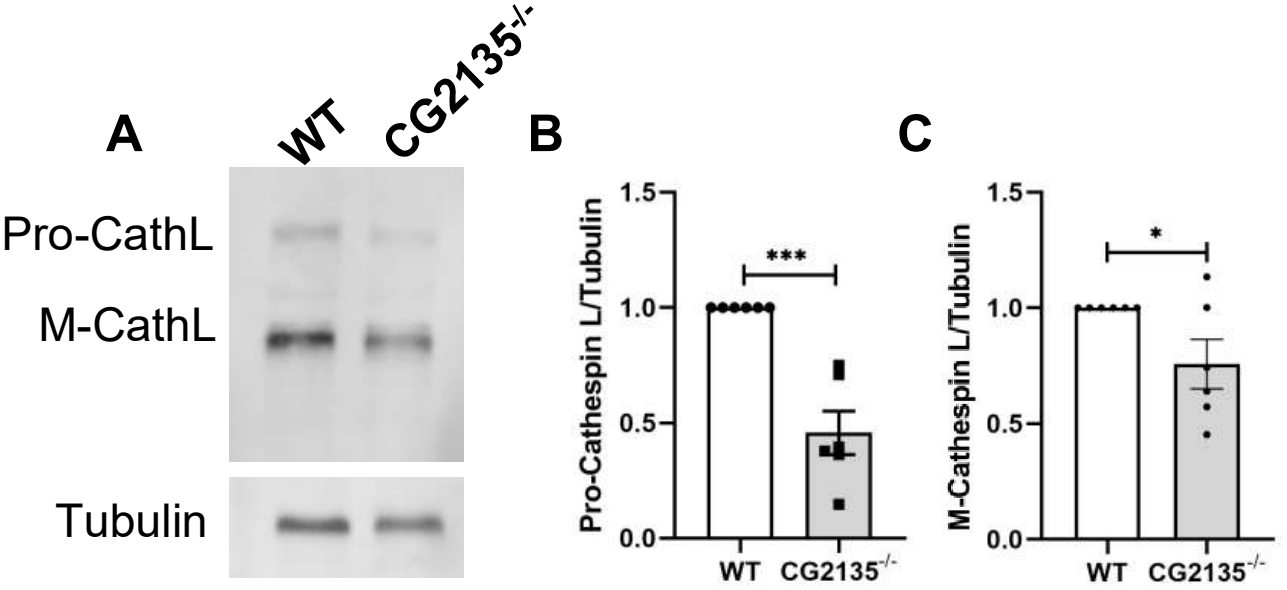
(A) Western blot of Cathepsin-L showing the protein level of Pro-CathL and Mature-CathL (37-25 kDa) from brain lysate of 30-day-old WT and CG2135^−/−^, as indicated in the figure. α-Tubulin (54 kDa) was used as a loading control. (B-C) The graph represents the quantification of Pro-Cathepsin-L/tubulin and Mature-Cathepsin-L/tubulin band intensity, showing a decreased level in the CG2135^−/−^ brain lysate. N = 6. Error bars represent the standard error of the mean (SEM). Asterisks represent a level of significance (***P ≤ 0.001, *P ≤ 0.05; Student’s t-test).

**Figure S3:**
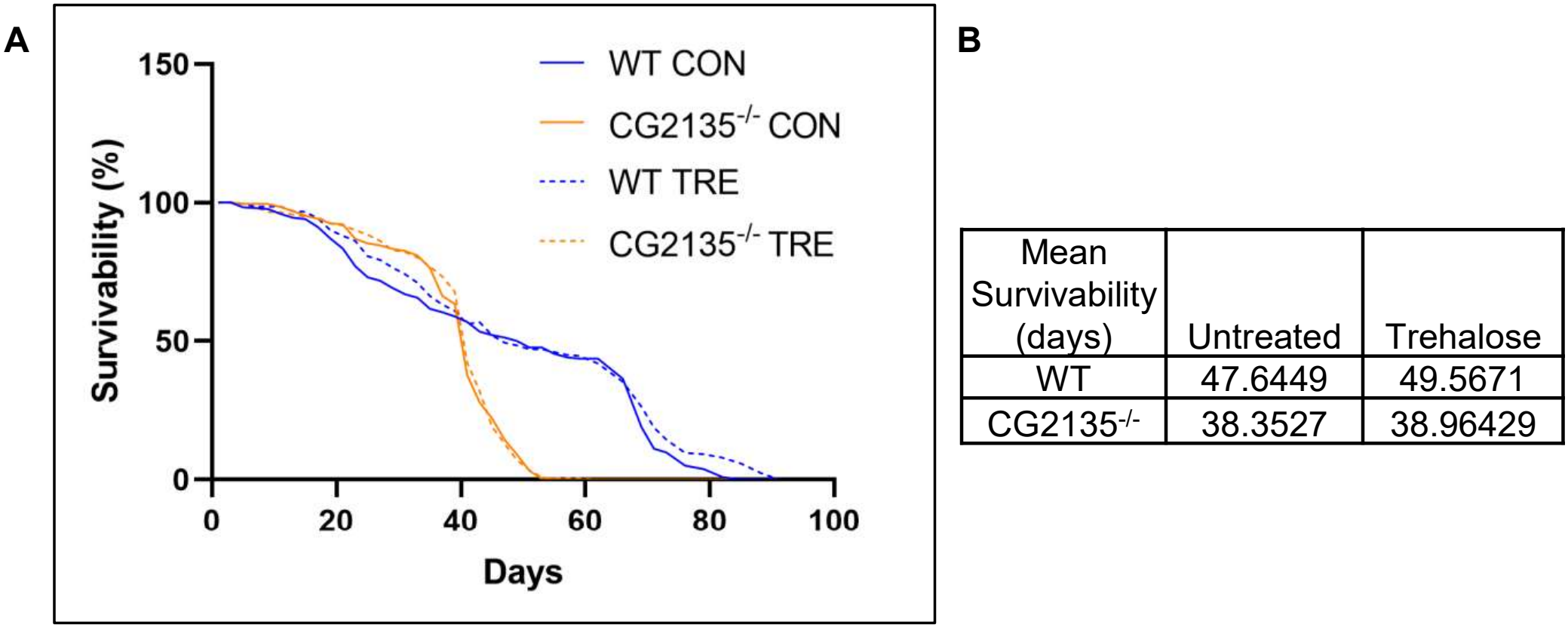
(A) Survival curve of the WT, CG2135^−/−^, and trehalose-treated WT and CG2135^−/−^ flies. (B) Mean survivability (in days) values are provided in the table below. N = 200 flies.

**Table S1:**
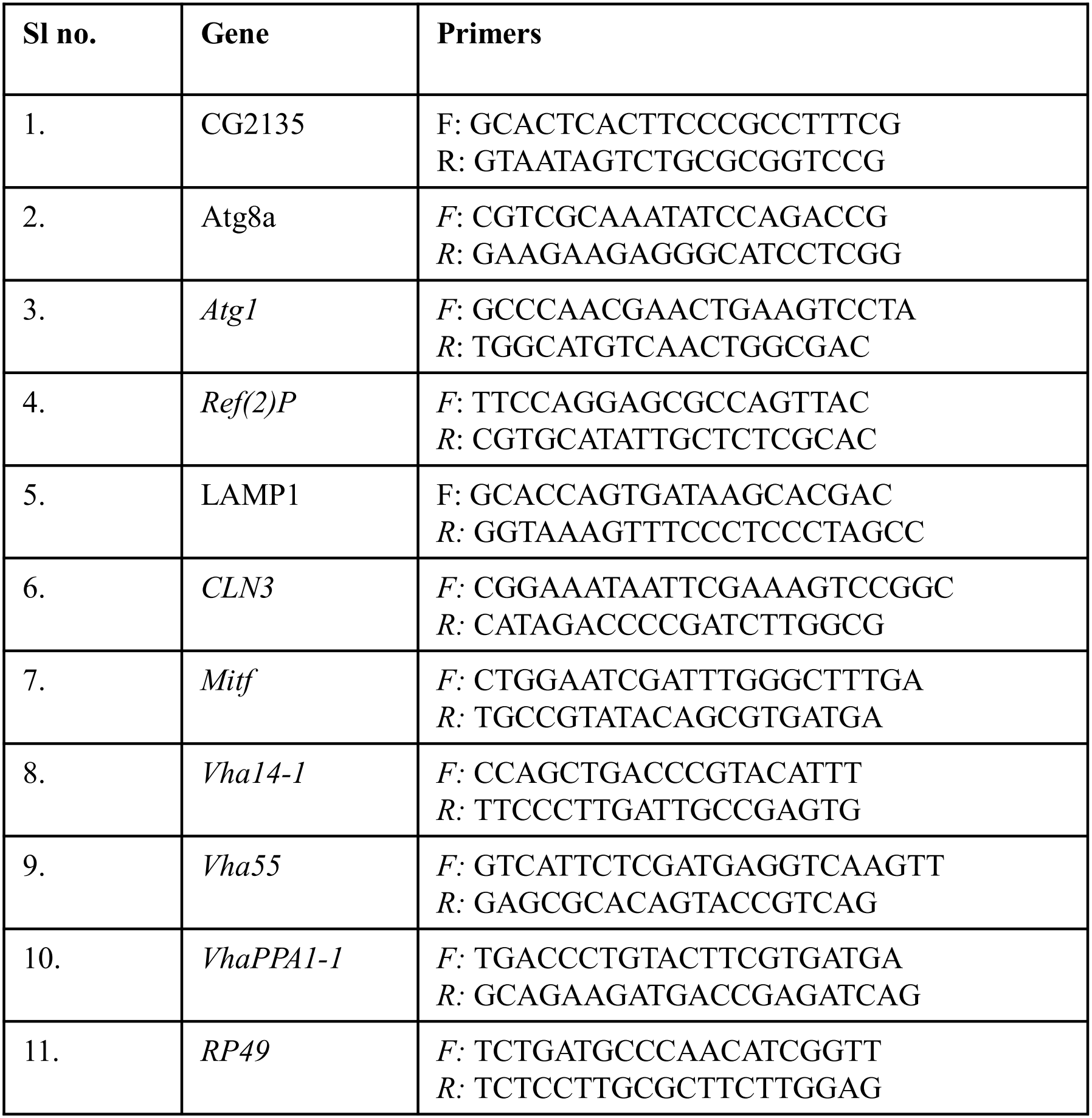
List of primers used in the study.

